# Inference of marker genes of subtle cell state changes via iterative logistic regression

**DOI:** 10.1101/2025.05.23.655858

**Authors:** Yingtong Liu, Aaron G. Baugh, Evanthia T. Roussos Torres, Adam L. MacLean

## Abstract

We present iterative logistic regression (iLR) for the identification of small sets of informative marker genes. Differential expression and marker gene selection methods for single-cell RNA sequencing (scRNAseq) data can struggle to identify small sets of informative genes, especially for subtle differences between cell states, as can be induced by disease or treatment. iLR applied logistic regression iteratively with a Pareto front optimization to balance gene set size with classification performance. We benchmark iLR on *in silico* datasets demonstrating comparable performance to the state-of-the-art at single-cell classification using only a fraction of the genes. We test iLR on its ability to distinguish neuronal cell subtypes in healthy vs. autism spectrum disorder patients and find it achieves high accuracy with small sets of disease-relevant genes. We apply iLR to investigate immunotherapeutic effects in cell types from different tumor microenvironments and find that iLR infers informative genes that translate across organs and even species (mouse-to-human) comparison. We predicted via iLR that entinostat acts in part through the modulation of myeloid cell differentiation routes in the lung microenvironment. Overall, iLR provides means to infer interpretable transcriptional signatures from complex datasets with prognostic or therapeutic potential.

## 1 INTRODUCTION

Single-cell resolution has proven essential for resolving certain cell states and their corresponding gene expression differences in biological systems. Single-cell RNA sequencing (scRNA-seq) quantifies gene expression at the single cell level, and with decreasing costs and a proliferation of protocols, has been widely adopted (Svensson et al., 2020). Downstream of cell clustering to identify cell states is the identification of marker genes that distinguish cells by their type, state, condition, etc (Huang & Zhang, 2021; Pullin & McCarthy, 2024). Marker genes are critical for interpreting cell clusters as different cell states. They are also essential for comparative analyses of cells across treatments, genetic perturbations or disease states (Dawson & Kouzarides, 2012; Falkenberg & Johnstone, 2014; Perri et al., 2017). When such comparisons reveal subtle transcriptional differences between groups, they can be difficult to capture with conventional marker gene selection approaches. For example, some epigenetic therapies induce broad shifts in gene expression without affecting target genes directly (Falkenberg & Johnstone, 2014). Even though certain epigenetic modulators have been approved and are being used as cancer therapeutics, their precise mechanisms of action often remain unclear (Dawson & Kouzarides, 2012; Perri et al., 2017). Furthermore, combination therapies—designed to enhance efficacy and prevent drug resistance—are increasingly common in clinical practice (Fitzgerald et al., 2006). These regimens often produce complex and context-dependent transcriptional responses, making it essential to identify condition-informative gene sets to elucidate therapeutic mechanisms and optimize treatment strategies.

A key challenge in the analysis of single-cell datasets to perturbations due to e.g. cancer or developmental disorders (Chea et al., 2023) is the identification of informative gene sets that capture the (often widespread bu subtle) transcriptional differences that occur. Canonical statistical methods for differential expression testing, such as DESeq2 (Anders & Huber, 2010), edgeR (Robinson et al., 2010), or the Student’s t or Wilcoxon rank sum tests are commonly applied (Wilcoxon, 1945). These approaches rely on ranking genes by statistical significance and it can be difficult to select compact, interpretable gene sets—particularly when changes are small in magnitude but biologically meaningful.

A number of methods have been developed specifically for marker gene selection in the context of cell state identification (Dumitrascu et al., 2021; Nelson et al., 2022; Sun & Qiu, 2024; Vargo & Gilbert, 2020). These approaches are effective at identifying genes that distinguish well-separated clusters, thereby facilitating cell type or state annotation (Pullin & McCarthy, 2024), but they are not always well-suited for detecting markers when the perturbation- or treatment-induced expression changes are subtle, widespread, and may not tightly aligned with cell state boundaries. Moreover, few methods provide principled means to control the size and the redundancy of the gene set, which can be critical for interpretability and downstream validation. Logistic regression is a simple and interpretable method for classification (Berkson, 1944; Cox, 1958). The coefficients of a fitted logistic regression model provide means for ranking features by importance, enabling feature selection. Logistic regression has demonstrated competitive performance in scRNA-seq classification (Huang & Zhang, 2021; Le et al., 2022) and marker gene selection (Pullin & McCarthy, 2024).

We developed iterative logistic regression (iLR) to combine cell classification and feature selection by iteratively refining a set of features via logistic regression. We used Pareto front optimization to balance the gene set size with classification accuracy, enabling the inference of small, informative gene sets. iLR is designed to capture condition- or treatment-driven differences, providing interpretable signatures while maintaining high classification accuracy—even for weak signals with high biological variability.

The remainder of this paper is organized as follows. In the next section we discuss the methods and implementation of iLR. We go on to assess the performance of iLR on *in silico* datasets against standard and state-of-the-art feature selection and classification methods. We tested iLR on a large human scRNA-seq dataset comprising multiple neuronal samples from autism patients and healthy controls (Velmeshev et al., 2019) and demonstrated that iLR can accurately distinguish differences with small sets of informative genes. We applied iLR to immune cell types in breast and lung metastatic tumor microenvironments (TMEs) to identify genes that mark for changes due to combination therapy (Roussos Torres et al., 2024; Sidiropoulos et al., 2022). In combination with downstream analysis via gene regulatory network inference, we identified putative mechanisms of action of combination therapy in immune cell subtypes.

## 2 METHODS

### 2.1 Overview of iLR

Single-cell RNA sequencing (scRNA-seq) datasets were split into training and testing sets for each evaluation. A logistic regression classifier is applied on the training set with n cells and m genes iteratively. At each iteration, we have fitted the logistic regression model:

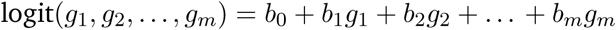

which is regularized by an L2 penalty with a rate of 0.1 to avoid overfitting. Coefficient *b_g_* for gene *g* as a measure of gene *g*’s significance in logistic regression classifier. Genes are ranked by coefficients and the top 80% genes are kept for the next iteration of logistic regression (Figure 1A). The algorithm proceeds until less than 10 genes are left. 5-fold cross-validation accuracy, AUC, and size of gene set are recorded at each iteration. iLR was implemented in Python and `LogisticRegression` with the `liblinear` solver from `scikit-learn` (Pedregosa et al., 2011). iLR uses L2 regularization with C=0.1.

**Figure 1:**
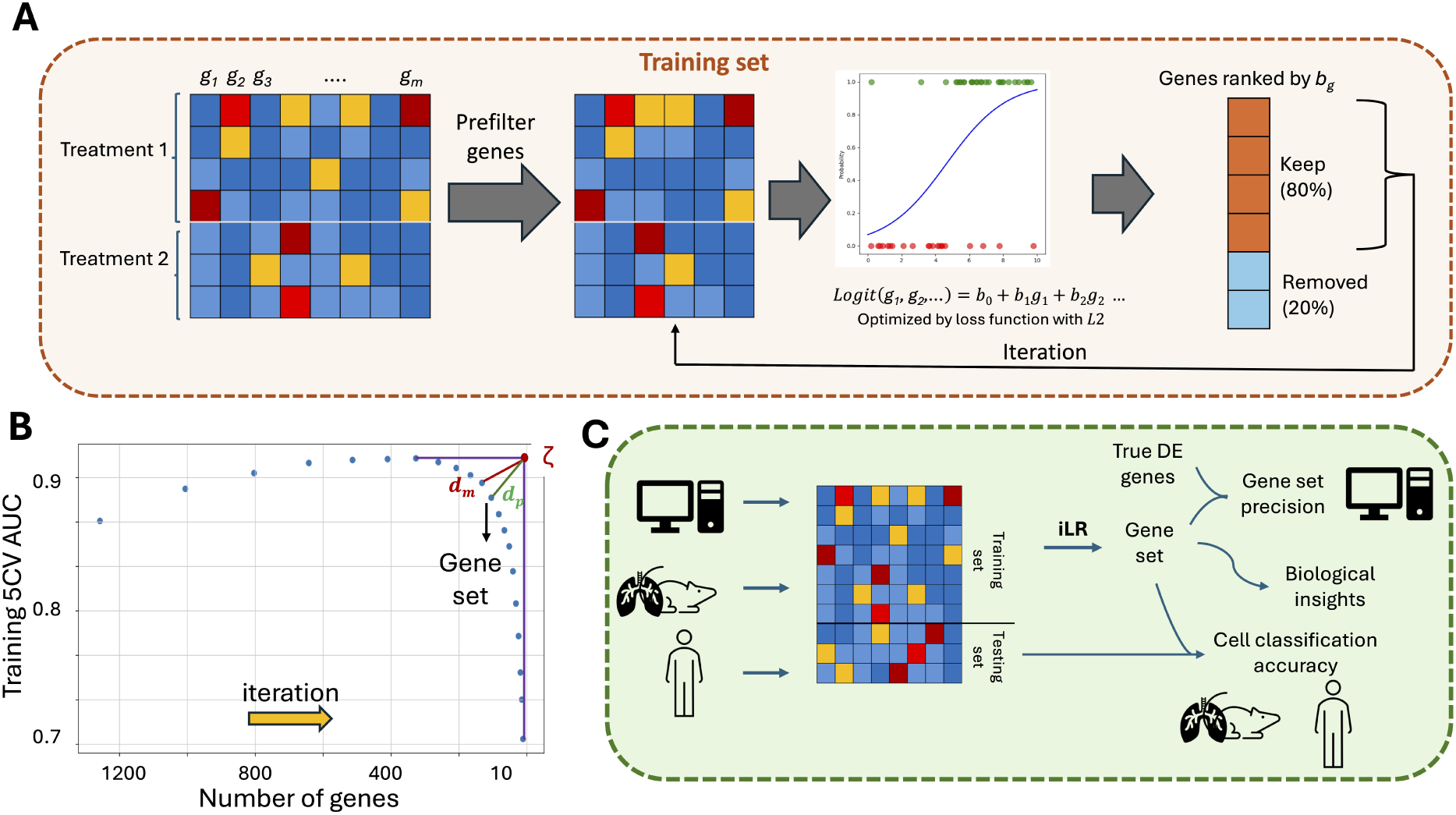
Overview of iLR. (**A**) The pipeline employed by iLR to select genes based on iterative logistic regression. (**B**) A Pareto front approach is taken to selection the optimal gene set. Each point indicates one gene set: over successive iterations the gene set size decreases. ζ marks the point of max. AUC and min. gene number from which the Pareto optimum is found via d_m_, the point that minimizes the distance to ζ among possible gene sets. In cases where a penalty is applied, d_m_ is replaced with d_p_, the penalized Pareto optimal gene set. (**C**) Training, testing, and applications of iLR are performed on simulated, murine, and human scRNA-seq datasets. Gene sets are evaluated against the ground truth, the literature, or comparatively across datasets from different organs or species.

### 2.2 Gene set size selection by Pareto front analysis

The gene set is selected based on the area under the curve (AUC) and the gene set size. The goal is to balance the classification AUC and gene set size to find relatively small sets of genes with the ability to classify cells correctly. A Pareto front is used for this task. The Pareto front provides means for multiobjective optimization: given the goal of finding a gene set with small size and high classification AUC, the AUC and the gene set size are normalized to a range between 0 and 1, and the intersection (*ζ*) between the highest AUC and lowest gene set size is found (Figure 1B). The optimal point is the one with minimal distance *d_m_* to *ζ* (Figure 1B). To adapt the criterion to allow for finding a smaller gene set at the expense of accuracy, we apply a user-defined penalty (*E*) to the distance such that a point with distance *d_p_* greater than *d_m_* to the intersection is selected (Figure 1B). To get stabilized gene sets, 10 rounds of iLR were done and genes appearing more than 5 times are in the final gene set.

### 2.3 Simulating scRNA sequencing data with Splatter

To test the iLR, we utilized the R package Splatter (Zappia et al., 2017) to simulate scRNA-seq data. There exist significant challenges in the analysis of methods using simulated scRNA-seq data (Crowell et al., 2023), nonetheless it remains a useful tool in combination with other analyses. Splatter allows the estimation of parameters from scRNA-seq data and the manipulation of the extent to which genes are differentially expressed. As an input sample, we used myeloid-derived suppressor cells (G-MDSCs) from untreated muring lung metastases, with 1202 genes. We varied two Splatter parameters to generate a total of eight datasets: the number of cells in the sample and the `DEscalefactor`, which was varied from 0.1 to 1 to capture very small to very large log fold change differences in gene expression (Figure 2A). The sample size was tested by varying the number of cells from either 500 or 2000. Each simulated dataset was balanced, i.e. 50% of cells in each condition (control and perturbed). The precision of the inferred gene set was calculated based on the known differentially expressed genes, and was used as a metric to assess the performance of iLR (Figure 1C). Splatter-generated raw count matrices were normalized to 10^4^ counts per cell and then log-transformed.

**Figure 2:**
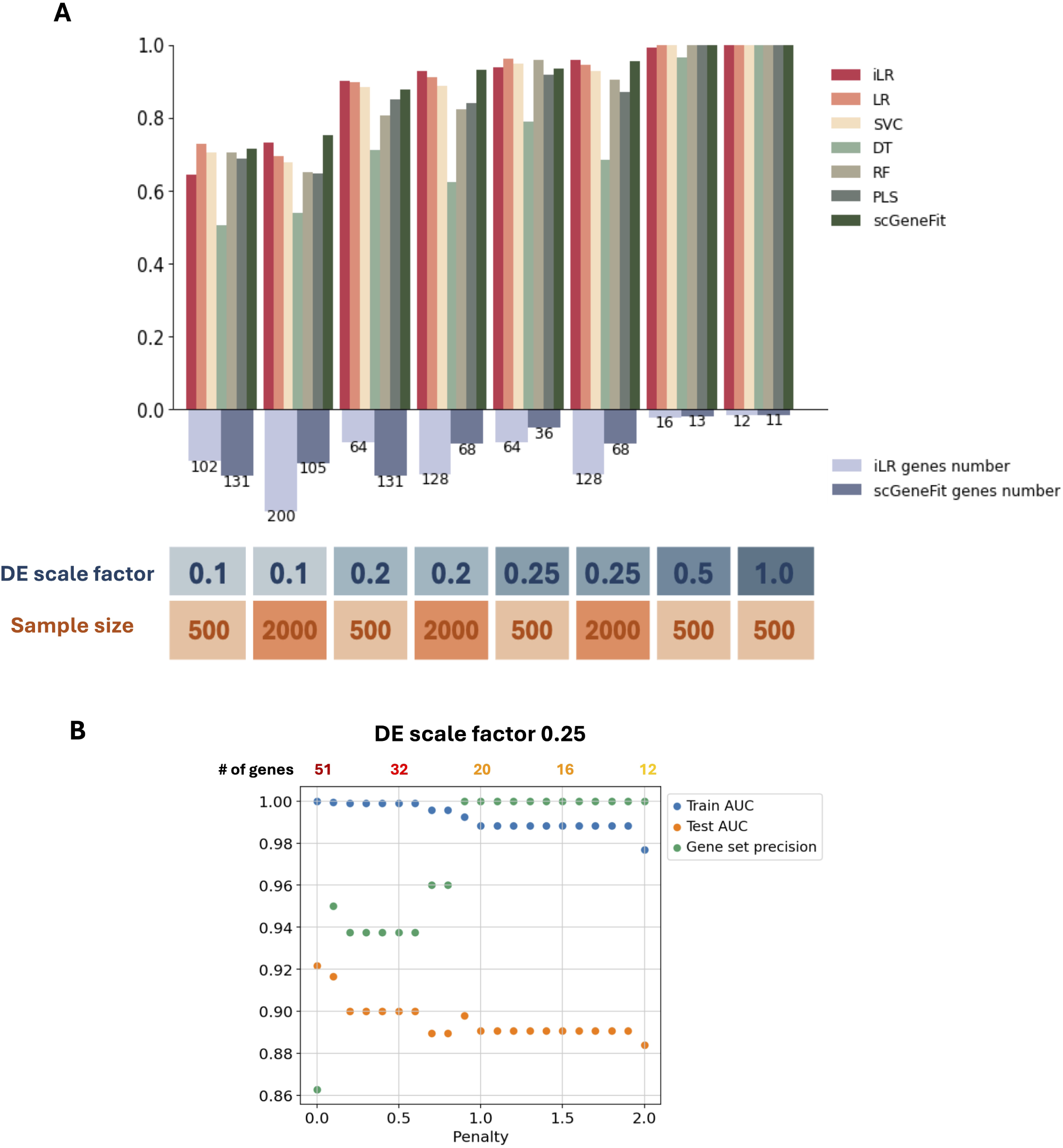
Benchmarking iLR on simulated datasets. (**A**) Comparison of iLR with other methods for classification via the AUC on simulated scRNA-seq datasets with different differential expression (DE) scale factors and sample sizes. The number of genes applies only to iLR and scGeneFit combined with pareto front without penalty. SVC: support vector classifier; DT: decision tree; RF: random forest; PLS: partial least squares. (**B**) The impact of the Pareto front penalty at scale factor 0.25 on the training & test AUC and the gene set precision.

### 2.4 Benchmarking iLR against alternative classification approaches and marker gene selection methods

To evaluate the capability of iLR in classification, several basic machine learning classification algorithms were applied to the Splatter simulated scRNA-seq data. The following algorithms were compared in performance to iLR (all implemented in `scikit-learn`). Simple logistic regression was implemented, regularized by an L2 penalty with rate 0.1. A linear support vector classification was implemented with a hinge loss. A decision tree was implemented with `min_sample_split` 10. A random forest was implemented with default parameters, and a partial least squares regression model was implemented with 12 components.

To assess the ability of iLR to accurately identify differentially expressed genes, we compared iLR genes to genes identified using a Wilcoxon rank sum test or scGeneFit (Dumitrascu et al., 2021) on simulated scRNA-seq data. scGeneFit utilizes the idea of compressive classification and largest margin nearest neighbor algorithm to select marker genes for cell states (Dumitrascu et al., 2021). To make them comparable, we fixed the gene set size to match the size we got from iLR. We selected the top genes in Wilcoxon rank sum test. Then we evaluated the gene sets derived from three methods by comparing them with the true DE genes and their capability of classification. Separately, we applied iLR, Wilcoxon sum rank test, and scGeneFit to NT2.5-LM scRNA-seq data to find the genes contributing to the treatment effects.

### 2.5 scGeneFit Gene Set Selection

Similar to iLR, we optimized scGeneFit gene sets using a Pareto front approach. In each round of scGeneFit we set the number of genes to approximately 80% of the previous round to fairly compare similarly sized gene sets. We also applied a Pareto front approach to the results of multiple runs of scGeneFit, balancing classification AUC and gene set size to infer the optimal number of genes to retain.

### 2.6 scRNA-seq data processing and analysis

#### 2.6.1 snRNA-seq from ASD patients and controls

Neuronal single-nuclei RNA sequencing (snRNA-seq) data from autism-spectrum disorder (ASD) patients and controls was analyzed (Velmeshev et al., 2019). Pre-processed data with cluster annotations is available from the UCSC Cell Browser at: https://autism.cells.ucsc.edu.

#### 2.6.2 Lung metastasis scRNA-seq library preparation and sequencing

Primary tumor scRNA sequencing data derived from mice injected with NT2.5 cell line with vehicle (V), entinostat (E), entinostat with aCTLA4 and aPD1 (EPC), entinostat with aCTLA4 (EC), and entinostat with aPD1 (EP) treated. For library preparation, 10x Genomics Chromium Single-Cell 3’ RNA-seq kits v2 were used. Gene expression libraries were prepared according to the manufacturer’s protocol. RNA was extracted from 20 whole tumors from the following groups: Vehicle control (V), entinostat-treated (E), entinostat and anti-PD-1 (EP), entinostat and anti-CTLA-4 (EC), entinostat with anti-PD-1 and anti-CTLA-4 (EPC), with four biological replicates from each of the five experimental groups. Tumors were sequenced in four batches: RunA (eight tumors; 1 E, 2 EP, 2 EC, 2 EPC, 1 V), RunB (eight tumors; 1 E, 2 EP, 2 EC, 2 EPC, 1 V), Pilot1 (two tumors, 1 E and 1 V), and Pilot2 (two tumors, 1 E and 1 V). Each batch had an approximately equal assortment of samples from each treatment group to reduce technical biases. Illumina HiSeqX Ten or NovaSeq were used to generate approximately 6.5 billion total reads.

#### 2.6.3 Alignment and data preprocessing

Paired-end reads were processed using CellRanger v3.0.2 and mapped to the mm10 transcriptome v1.2.0 by 10× Genomics with default settings. ScanPy v1.9.1 (Wolf et al., 2018) and Python v3 were used for quality control and basic filtering. For gene filtering, all genes expressed in less than 3 cells within a tumor were removed. Cells expressing less than 200 genes or more than 8,000 genes or having more than 15% mitochondrial gene expression were also removed. Gene expression was total count normalized to 10,000 reads per cell and log-transformed. Highly variable genes were identified using default `ScanPy` parameters, and the total counts per cell and the percent mitochondrial genes expressed were regressed out. Finally, gene expression was scaled to unit variance, and values exceeding 10 standard deviations were removed. There were 54,636 cells and 19,606 genes after pre-processing. Batch effects were corrected using the `ComBat` batch correction package. Neighborhood graphs were constructed using 10 nearest neighbors and 30 principal components. Tumors were clustered together using Louvain clustering and 6 main clusters were identified (with resolution parameter 0.1).

#### 2.6.4 Application of iLR to lung metastatic and primary tumor samples

We analyzed differences in each TME due to two treatments: entinostat alone (E) or in combination with anti-PD-1 and anti-CTLA4 (EPC) against vehicle (V), because 1) we were most interested in the mechanisms underlying these effects, and 2) these treatments were shared across two TME datasets (NT2.5 in the primary breast and NT2.5LM in the lunch). This enabled comparison of iLE gene sets across organs. E and EPC are also the most clinically relevant and have been evaluated in clinical trials (Roussos Torres et al., 2024).

Given the inherent differences in cell states and gene expression profiles between the two tumor models, we adjusted the normalized count data for cancer cells and mature myeloids. To better isolate genes relevant to treatment effects in cancer cells and avoid confounding by metastatic signatures we first identified genes that were differentially expressed between two metastatic breast cancer cell lines (NT2.5LM and its parental line NT2.5 (Baugh, Gonzalez, Narumi, Kreger, Liu, Rafie, Castanon, Jang, Kagohara, Anastasiadou, Leatherman, Armstrong, Chan, Karagiannis, Jaffee, MacLean, & Roussos Torres, 2024). These genes were filtered out prior to applying iLR. We used classification AUC from lung metastasis-trained models tested on primary tumors to determine the optimal number of genes to exclude for each treatment comparison. Specifically, the top 1,000 genes were removed in the V vs. E comparison, and the top 500 in the V vs. EPC comparison. For mature myeloid cells, dendritic cells were excluded from the lung metastasis samples. For each cell type, we applied `rank_gene_groups` with the Wilcoxon rank-sum test to identify significantly differentially expressed genes to be used as input for iLR.

### 2.7 Gene regulatory network inference based on marker gene sets

To investigate genes underlying treatment effects (V vs E and V vs EPC) in breast cancer, we applied SCORPION (Osorio et al., 2024b) using gene sets identified by iLR and scGeneFit with Pareto front penalty 0. We used TF-target database Dorothea (Badia-i-Mompel et al., 2022; Garcia-Alonso et al., 2019; Müller-Dott et al., 2023) and protein-protein interaction database STRING (Szklarczyk et al., 2023). SCORPION was run with default parameters. Comparisons included iLR and scGeneFit gene sets under both treatment conditions. SCORPION was run with default parameters, and low-weight edges in the TF–target gene network were filtered to reduce noise. Using the vehicle group as baseline, we removed the bottom 50% of edges by weight, which accounted for only about 12% of the total network weight. The same cutoff was applied to the treatment groups for consistency.

To identify transcription factors whose regulatory roles were significantly altered by treatment, we performed Wilcoxon tests separately on activating and repressing interactions. Volcano plots were used to highlight TFs with significant changes in regulatory activity.

### 2.8 Statistics

Hypergeometric tests were used to assess the significance of the overlap of two gene lists. iLR gene lists from human ASD and control snRNA sequencing data were evaluated by overlapping with all the genes in the SFARI Gene Module database (Abrahams et al., 2013), which contains genes having evidence of genetic association with ASD. A Bonferroni correction was applied to account for multiple testing. Biological processes were analyzed via gene ontology (GO) analysis (Feuermann et al., 2025) in the Python package `GSEApy` (Fang et al., 2023).

## 3 RESULTS

### 3.1 iLR classifies single cells accurately with small sets of genes from simulated scRNA-seq data

To test the ability of iLR to classify cells from scRNA-seq data and identify useful gene sets, it was applied to eight scRNA-seq datasets simulated via Splatter (Zappia et al., 2017) with different scales of log fold change of gene expression between conditions and different sample sizes. The gene set is selected based on the AUC via the Pareto front (Figure 1). Overall, we found that iLR outperformed alternative methods for cell classification, especially with relatively small scale factors (Figure 2A, Figure S1A). Moreover, iLR classified cells accurately with many fewer genes (1% to 16% of the total genes) than alternative approaches. scGeneFit applied with a Pareto front (as in iLR) performed similarly in terms of classification accuracy across different DE scales (Figure 2A). At the smallest scale factors, 0.1, we found that increasing the sample size from 500 to 2000 could improve classification accuracy; here iLR selected larger sets of genes. Overall, larger sample sizes or smaller scale factors required larger gene set sizes to achieve high AUC.

We found that applying a penalty to the Pareto from optimization to favor smaller gene set sizes over highest possible AUC allowed iLR to identify smaller gene sets without significant loss in classification accuracy (Figure 2B). Notably, at a scale factor of 0.25, applying Pareto front analysis with a penalty *≥* 0.9 was able to identify a small, informative gene sets containing all the true differentially expressed (DE) genes.

Wilcoxon rank sum tests can often identify good candidate sets of marker genes (Pullin & McCarthy, 2024), but they are not designed to control for the number of significant genes. As such, applied to Splatter datasets the number of identified genes at a p-value cutoff of 0.05 varied widely from 4 to 222 (Table 1): the number of DE genes increases with the sample size and the scale factor. This can be mitigated by incorporating a Pareto front approach with penalty constraints. For instance, with a penalty of 2, the gene set size was limited to around 20 genes (Figure S1A).

**Table 1:**
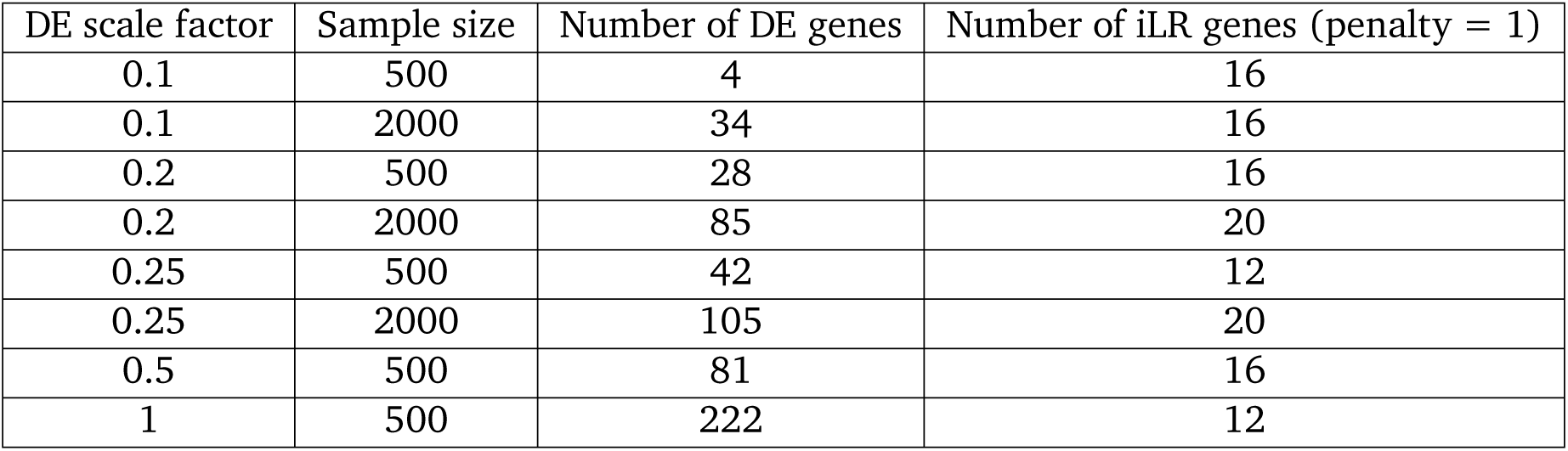
The number of significant differentially expressed genes (adjusted p-value cutoff 0.05) identified by Wilcoxon rank sum test in eight simulated data.

Using Splatter datasets, we tested the robustness of iLR by running it ten times on the same dataset, varying the random seed. The number of genes selected remained consistent across all ten runs, although a slightly larger gene set size was selected in 3/10 runs, highlighting stochasticity in Pareto front optimization (Figure S1B).

### 3.2 Validation of iLR on snRNA-seq of autism spectrum disorder patients

To evaluate the performance of iLR to distinguish biological differences in human cell types, iLR was applied to a total of 17 cell states in the human brain to compare healthy vs. autism spectrum disorder (ASD) differences (Velmeshev et al., 2019)(Figure 3A). Three different criteria were used to select the optimal gene set: the gene set with the highest AUC (i.e. no Pareto front applied), the optimal gene set according to the Pareto front without penalty, and the optimal gene set according to the Pareto front with penalty 1 (Figure 3B, Figure S2A). Similar to the results obtained on simulated data (Figure 2B), applying a Pareto front with penalty 1 produced smaller gene sets while maintaining high accuracy (AUC values, (Figure 3B, **??**A). Among all cell states, L2/3, L5/6-CC, and L4 exhibited the highest AUC, indicating that these cell states exhibited the clearest differences between ASD patients and controls. This agrees with previous work on the relevance of L2/3, L5/6-CC, and IN-PV to ASD via high correlations between individual-level gene expression changes and ASD clinical severity scores (Velmeshev et al., 2019). In Velmeshev et al., microglia were predicted to have many changes in terms of gene expression from ASD to healthy controls (Velmeshev et al., 2019). here, iLR did not identify many genes associated with disease status nor classify the microglia accurately with a small gene set (Figure 3B). To investigate this, a Wilcoxon rank sum test was performed on the training samples from each patient and all control samples. The number of significant DE genes in microglia between ASD and controls was small, indicating that microglia changes resulting from ASD are heterogeneous and the majority of DE genes in microglia from ASD patients are unique to each patient (Figure S2B). In L2/3, in contrast, many DE genes were shared among patients (Figure S2C). This highlights the utility of iLR: in cases where classification accuracies are lower, there is likely more biological variability in the dataset worth of investigation, as is the case here with intrinsic heterogeneity of microglia.

**Figure 3:**
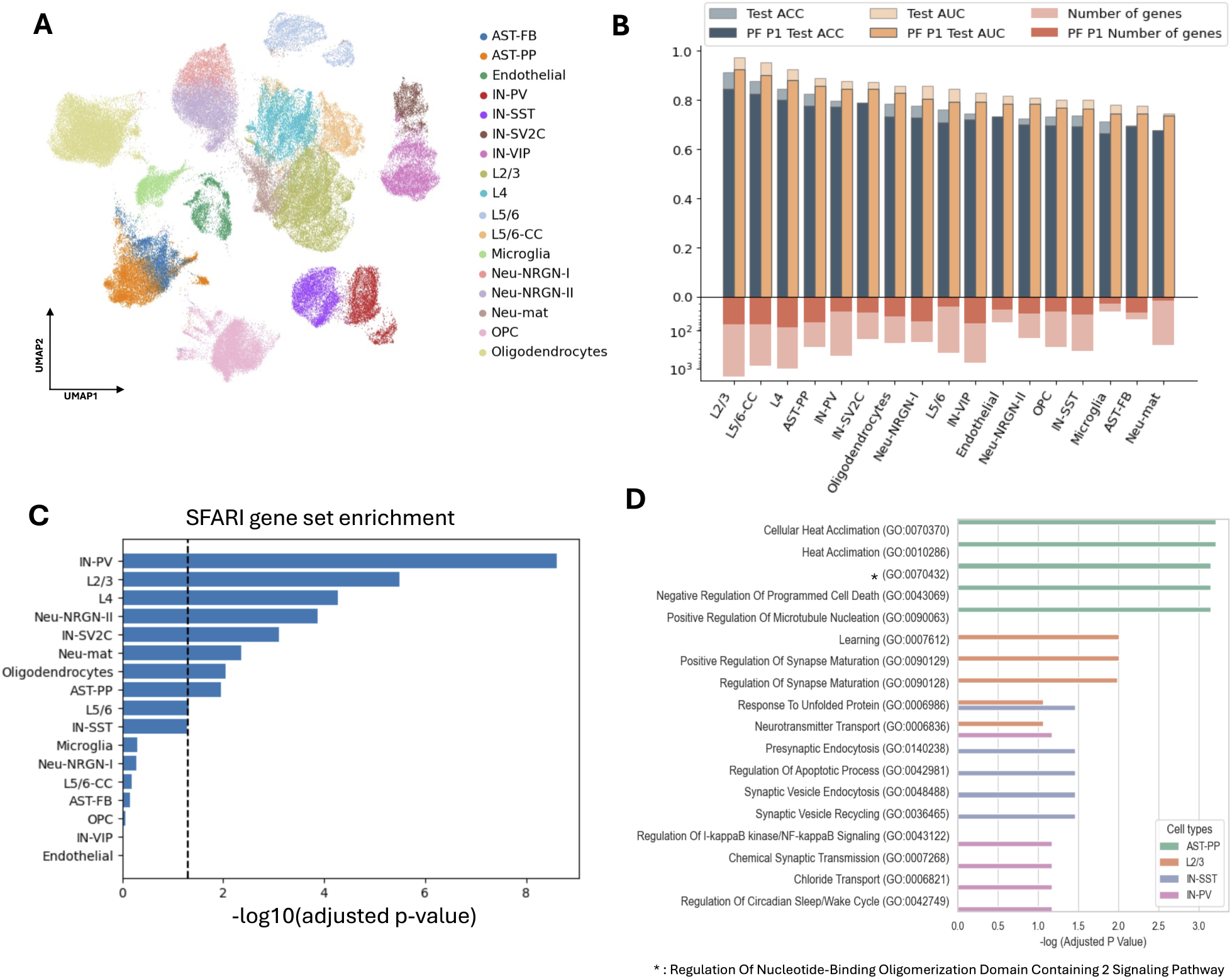
Assessment of iLR on neuronal cell fates in the brain. (**A**) UMAP depicting the 17 cell states identified in healthy control and autism spectrum disorder (ASD) in data from Velmeshev et al. (2019). (**B**) iLR classification accuracy (on ASD vs. control) and gene set size for each neuronal cell state. Comparison without (bold color) vs. with (pale color) a Pareto front penalty of 1 applied. (**C**) Enrichment in SFARI of the genes identified by iLR for each neuronal cell states. The dashed line denotes an adjusted p-value of 0.05. (**D**) Gene set enrichment analysis for genes identified in AST-PP, L2/3, IN-SST, and IN-PV cells.

To assess the genes identified by iLR, we calculated enrichment scores using the SFARI database (Abrahams et al., 2013), a comprehensive and validated repository of autism-associated genes. Using a hypergeometric test, we analyzed the enrichment of iLR-selected gene sets in SFARI for each cell state. The enrichment analysis revealed that IN-PV, L2/3, and L4 had the highest overlap with SFARI genes (Figure 3C), consistent with findings by Velmeshev et al.(Velmeshev et al., 2019) and Wamsley et al.(Wamsley et al., 2024). These results strongly support the conclusion that iLR can identify small sets of genes with high relevance, in this cases revealing differences between neuronal cell states in ASD vs control brains.

We conducted a Gene Ontology (GO) enrichment analysis for AST-PP, L2/3, IN-SST, and IN-PV. GO analysis revealed top terms related to brain and synaptic functions (Figure 3D). For IN-SST cells, synaptic regulation emerged as the most significantly altered term, consistent with (Wamsley et al., 2024). Terms associated with synaptic regulation, maturation, and neurotransmitter regulation were significantly enriched in L2/3, in agreement with (Wamsley et al., 2024); similar agreement was seen for AST-PP, enriching for apoptosis and cytoskeleton development. These findings collectively demonstrate the robustness and biological relevance of iLR for identifying gene sets in the context of ASD that can provide valuable insights into underlying molecular mechanisms.

### 3.3 iLR genes capture immunotherapy treatment effects by comparing changes in cell types across TMEs

We next applied iLR to two large scRNA-seq datasets describing cell types present in different tumor microenvironments (TMEs), where tumors are treated with immune checkpoint inhibitors combined with entinostat, a histone deacetylase inhibitor (Baugh, Gonzalez, Narumi, Kreger, Liu, Rafie, Castanon, Jang, Kagohara, Anastasiadou, Leatherman, Armstrong, Chan, Karagiannis, Jaffee, MacLean, & Torres, 2024; Sidiropoulos et al., 2022). With limited prior knowledge of the effects of treatment across cell types from different TMEs, we applied iLR to compare genes marking for treatment differences between the primary tumor (breast TME) and at metastatic sites in the lungs (lung TME). Using iLR, we analyzed MDSCs, T cells, mature myeloid cells, and the tumor across these TMEs (Figure 4A).

**Figure 4:**
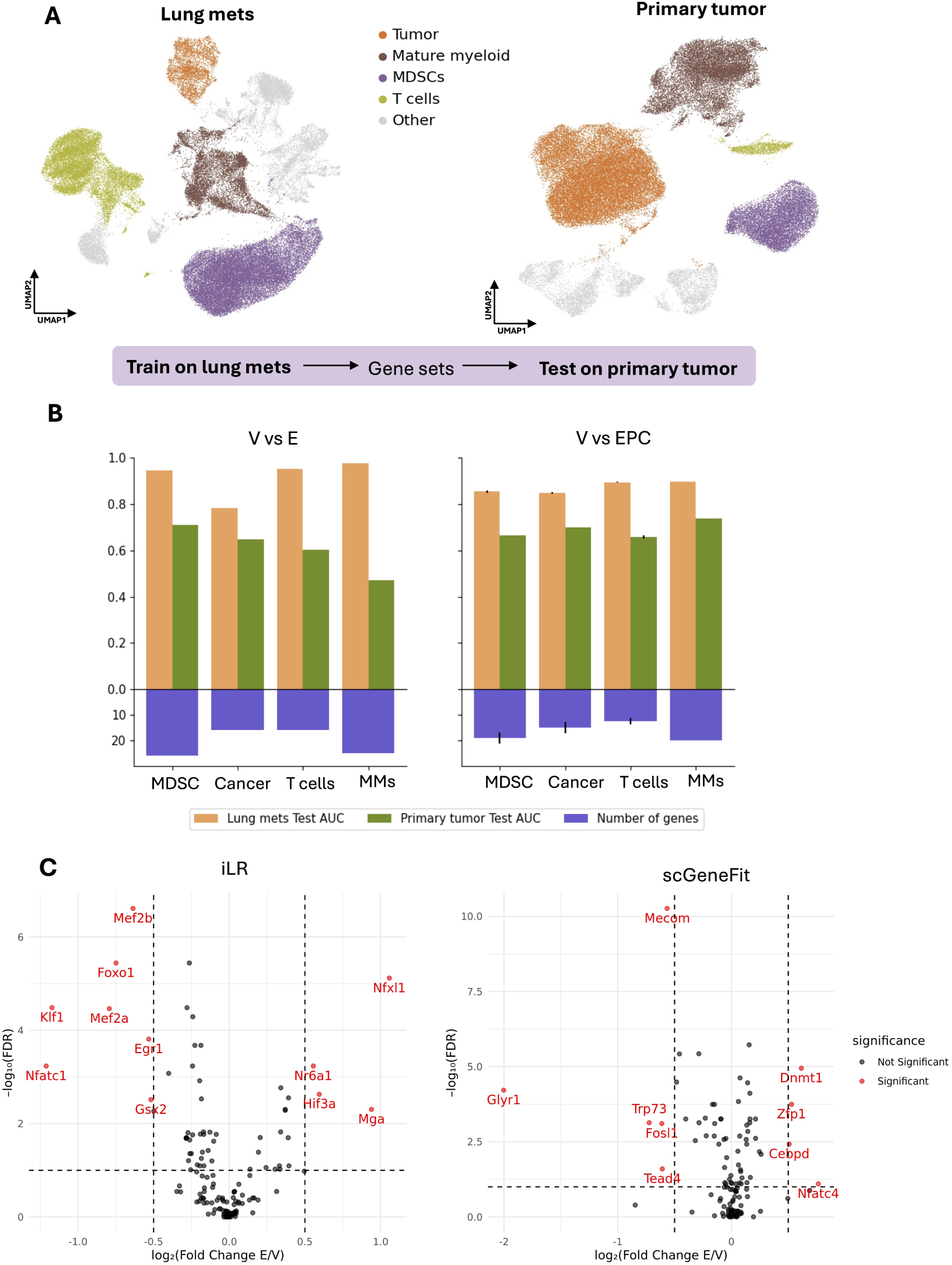
Analysis and discovery of the effects of immunotherapy on MDSCs. (**A**) Illustration of the high-level cell states that are shared between TME in the primary breast and the metastatic lung. (**B**) The accuracy of iLR in classification (via test AUC) for models trained on the lung mets and tested on the primary tumor. (**C**) Results of GRN inference in SCORPION: differential activity of Inhibitory TFs based on networks inferred using gene sets marking for the change with E or with EPC with respect to V, using iLR (left) or scGeneFit (right).

We first identified cell type-specific gene sets marking for treatment effects in the lung TME, and tested these for their ability to train a classifier for the same cell type in the breast. For changes induced by EPC treatment, all four cell states showed generally consistent AUCs tested on the primary breast TME (Figure 4B). This suggests that these gene sets identified based on lung metastatic differences can partially explain the effects of EPC in different TMEs. In contrast, comparison of V vs. E treatment showed variable results, where the mature myeloid population in the breast could not be classified using the lung mets iLR gene set (Figure 4B). We also performed the reciprocal analysis: identifying gene sets using iLR in the breast TME and testing them on cell state differences in the lung TME (Figure S3B). The results suggest a commutability in classification of cells based on iLR gene sets across these TMEs, indicating underlying similarities in transcriptional shifts with treatment. Notably, the gene set inferred from mature myeloid cells in the primary breast TME was more generalizable than that of the lung TME (Figure S3B). Direct comparisons of marker gene sets revealed limited overlap between TMEs (Figure S3A).

Going beyond unsupervised assessment of iLR gene sets, biologically several important genes were identified. In the mature myeloid cells from the lung TME, iLR identified two genes (out of an 18 gene set) associated with the effects of EPC that are highly relevant to cancer progression: *Vim* and *Ccr5*. Both these genes were ranked lower than 250*^th^* place by p-value using a Wilcoxon rank-sum test. Vim is a mesenchymal marker in tumor cells, and recent studies have shown that Vim-high macrophages promote tumor progression in hepatocellular carcinoma (Qiu et al., 2024). Ccr5 inhibition has been associated with anti-tumoral macrophage morphology (Jiao et al., 2019).

### 3.4 Analysis of iLR gene sets representative of differences in MDSCs yields insight into mechanisms of immunosuppression

Given the known role yet unclear mechanisms of MDSC immune suppression in breast cancer (Christmas et al., 2018; Kreger et al., 2023), we further analyzed gene sets characterizing differences in MDSCs with entinostat treatment. First, through comparison with gene expression differences observed in bulk RNA-seq from patient biopsies undergoing the same EPC treatment and in our mouse model, we found eight overlapping genes, highlighting conserved features across species. Of these, several stood out. Lcn2 was identified in both the murine scRNA-seq data (iLR gene set) and as a DE gene in clinical biopsies: Lcn2 is a well-established non-invasive diagnostic and prognostic marker for breast cancer progression (Hu et al., 2018). Another shared gene, Cd52, has been reported as a prognostic marker with the potential to predict the progression and stages of breast carcinoma (Ma et al., 2021). Similarly, Rps29, a ribosomal protein gene, is known to induce apoptosis and enhance the efficacy of anti-tumoral drugs (El Khoury & Nasr, 2021). In contrast, analysis of the comparable gene sets identified using scGeneFit found some overlap (Cd52), but overall fewer genes relevant to tumor progression, identifying many ribosomal genes which are less biologically informative. These findings suggest that iLR-identified genes from murine scRNA-seq can provide valuable insights with translational relevance.

To investigate possible mechanisms mediating treatment effects in MDSCs, we studied gene regulatory relationships predicted by gene sets inferred by iLR or scGeneFit. We inferred gene regulatory networks using SCORPION (Osorio et al., 2024a), which uses a message-passing algorithm and is specifically designed for the comparison of regulatory networks across conditions. We ran SCORPION using either the iLR or the scGeneFit gene sets as input and studied the resulting transcription factors (TFs) predicted to be significantly altered by treatment (Figure 4C). Of the top TFs predicted for input iLR genes, many were found to be involved in myeloid cell differentiation and MDSC suppressive function. *Egr1* restricts differentiation of myeloid cells to the (more suppressive) macrophage over the granulocyte lineage, as well as induce matrix metalloprotease 9 (MMP9) expression in MDSCs, which can increase invasion in the TME. *Foxo1* has been found to promote MDSC differentiation towards more mature myeloid cells; *Foxo1* deficiency can decrease MDSC suppressive function. Other TFs corroborate these predictions: *Nfatc1* shares targets with *Foxo1* and *Mef2a/b* in directing myeloid cell differentiation to affect MDSC fates.

Analysis of the TFs predicted to control treatment effects by SCORPION using scGeneFit genes as input found less evidence of TF regulation of myeloid cell fates. The presence of epigenetic regulators (*Dnmt1, Glyr1, Cepbd*) may in part be due to the effects on MDSCs of the epigenetic regulator entinostat.

## 4 DISCUSSION

We developed iLR to identify small and informative gene sets that explain treatment effects or disease conditions using scRNA-seq data, while also demonstrating strong classification accuracy. We validated iLR on simulated data and ASD patient datasets, and in application to the lung TME we demonstrated generalizability of gene sets. Key to the success of iLR was the use of Pareto front optimization to reduce the gene set size while maintaining accurate cell classification. Key therapeutic effect-associated genes included *Lcn2* and *Cd52*. GRN analysis highlighted transcription factors that regulated genes changing significantly with treatment.

iLR gene sets highlighted biological mechanisms of interest. Nonetheless, any gene set predicted via linear models is limited in its scope: downstream & nonlinear analyses of genes inferred by iLR or otherwise are important. These might include cell-cell communication network inference (Jin et al., 2021; Mitchell et al., 2025), GRN inference (Kamimoto et al., 2023; Rommelfanger et al., 2023; Wang et al., 2023), or generative modeling (Lopez et al., 2018; Mitra & MacLean, 2021). Transcription factors predicted through GRN inference that target adhesive or secreted proteins offer potential to bridge from cell-internal signaling pathways to extracellular signaling, as other methods have employed (Yan et al., 2025).

iLR-identified genes highlighted biologically relevant mechanisms in the cases of both the ASD and breast cancer analyses. For ASD, the genes highlighted by iLR closely aligned with the recent literature. In the case of breast cancer TMEs, genes identified by iLR prompted new hypotheses regarding mechanisms of immune suppression (via myeloid differentiation) and offered translational potential as they could mark for treatment effects across tumor contexts—from primary to metastatic site and even from mouse to human. ScGeneFit — especially when combined with Pareto front optimization as applied here — showed comparable performance at classification tasks and offers an alternative to iLR. However, in analyses of TMEs, scGeneFit identified gene sets consisting of many less biologically informative ribosomal genes. In downstream analyses of gene sets via GRN inference, scGeneFit identified well-known regulators whereas iLR identified putatively new regulators of immune function, potentially guiding discovery.

While iLR demonstrated strong predictive performance, it is inherently constrained by the assumption of linear relationships in logistic regression, potentially overlooking genes with nonlinear contributions. Additionally, despite the use of L2 regularization and feature elimination, multicollinearity among genes may still hinder the model’s ability to identify truly informative features, as correlated genes can mask each other’s effects. As illustrated in this work, downstream GRN analysis may help recover such missed genes by leveraging co-expression or regulatory patterns. Finally, like other computational approaches, iLR identifies associations rather than causality; experimental validation remains essential to confirm the functional relevance of selected genes.

Overall, iLR identified small, informative gene sets that captured subtle transcriptional cell state shifts, as can be often observed with developmental disorders, cancer treatment, or other therapeutic interventions. Analysis via both *in silico* datasets and real-world scRNA-seq data demonstrated that iLR can provide interpretable signatures while maintaining high classification accuracy. Coupled with downstream gene regulatory and other analyses, iLR can help guide the discovery of biological mechanisms & new biomarkers, and, in doing so, we can better understand the transcriptional basis of complex disease.

## 5 SUPPLEMENTARY TABLES and FIGURES

**Figure S1:**
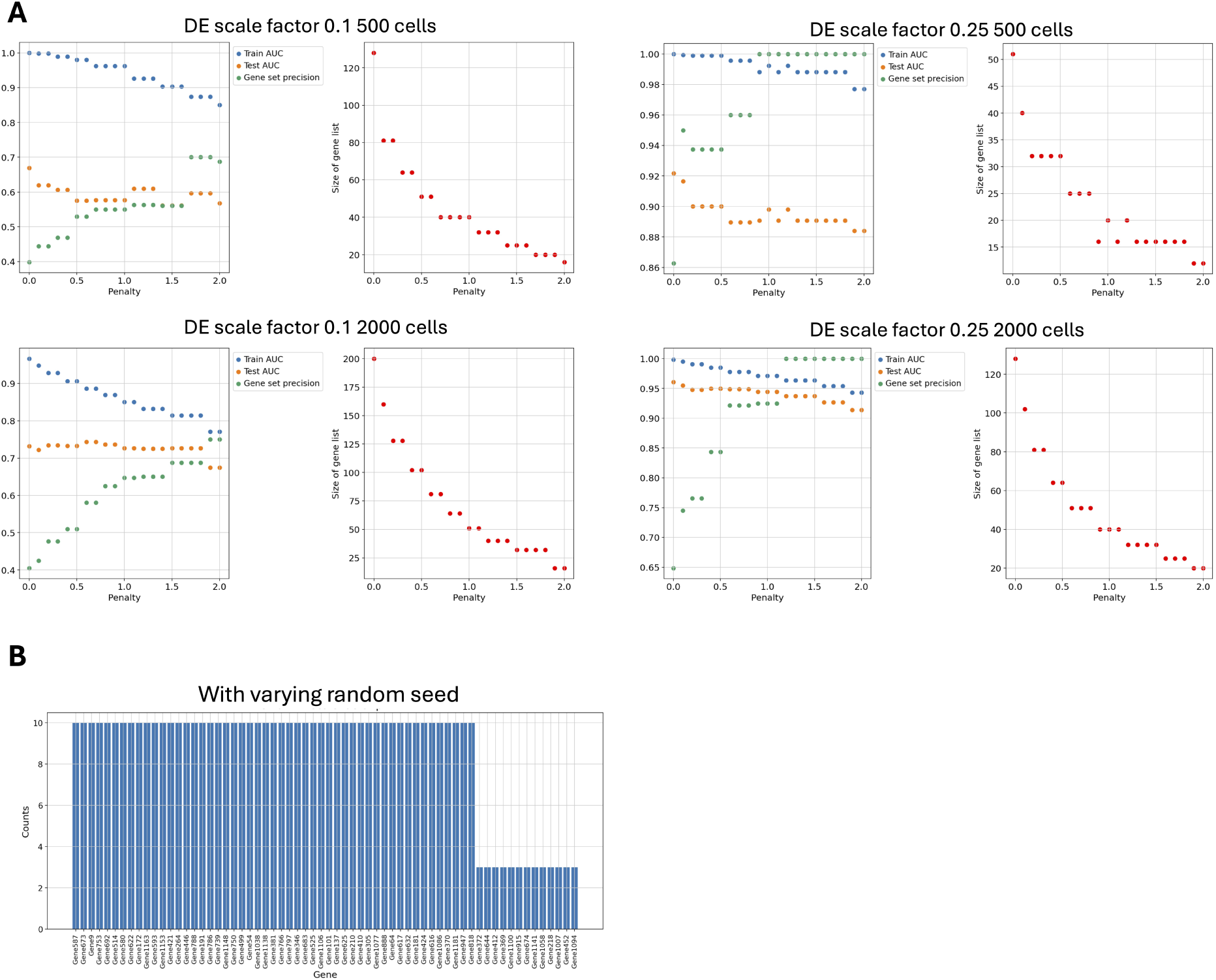
iLR evaluation on simulated data. (**A**) Scatter plots trace the penalty effect on four simulated data with different DE scale factor or sample size. (**B**) Examination the iLR robustness by running 10 times, varying the random seed.

**Figure S2:**
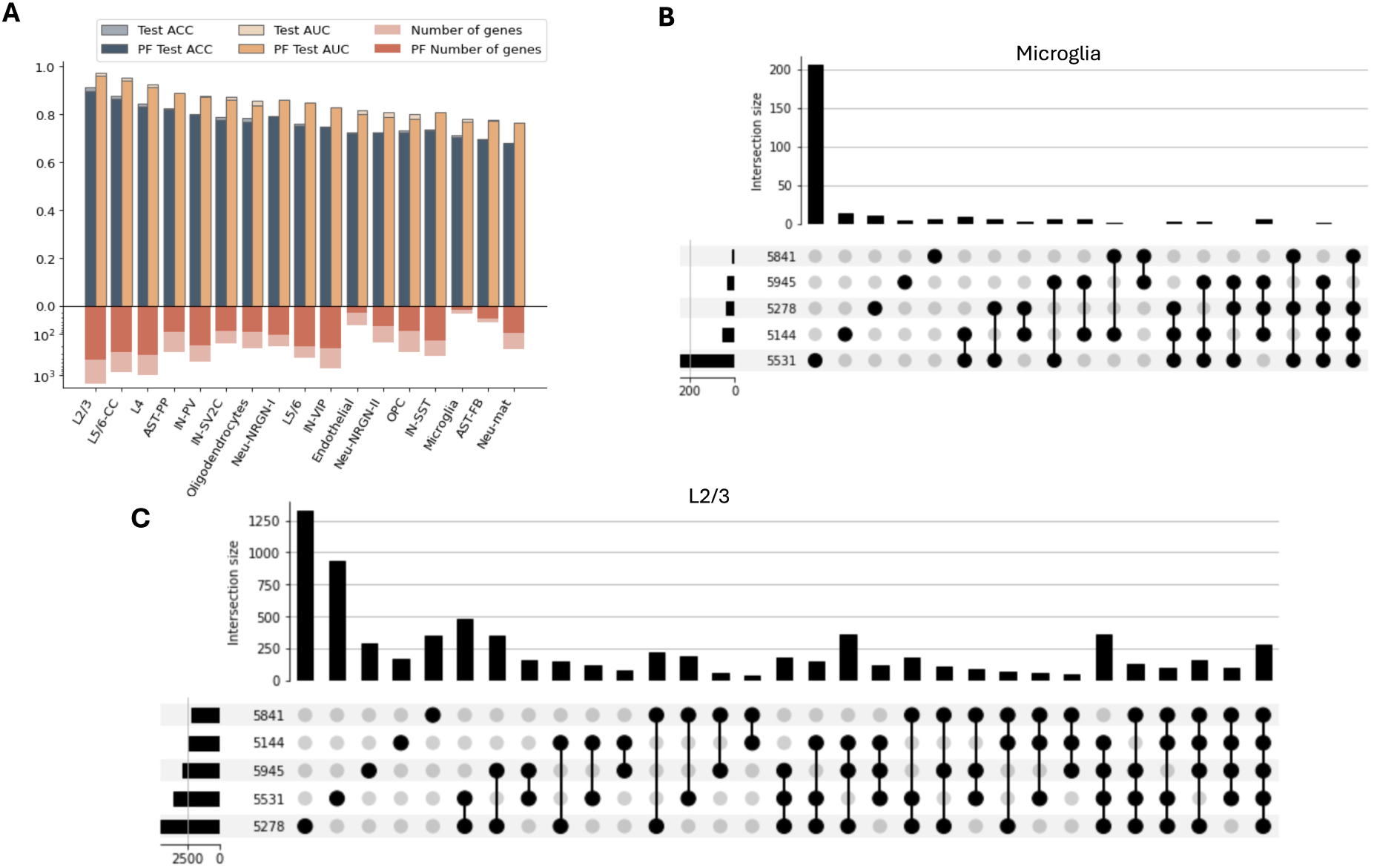
iLR application on ASD dataset. (**A**) iLR classification accuracy and gene set size comparing no pareto front (solid colored) and with pareto front without penalty (pale colored). (**B**) Upset plot of Wilcoxon rank sum test gene sets identified by comparison of microglia of individual patient and all controls. (**C**) Upset plot of Wilcoxon rank sum test gene sets identified by comparison of L2/3 of individual patient and all controls.

**Figure S3:**
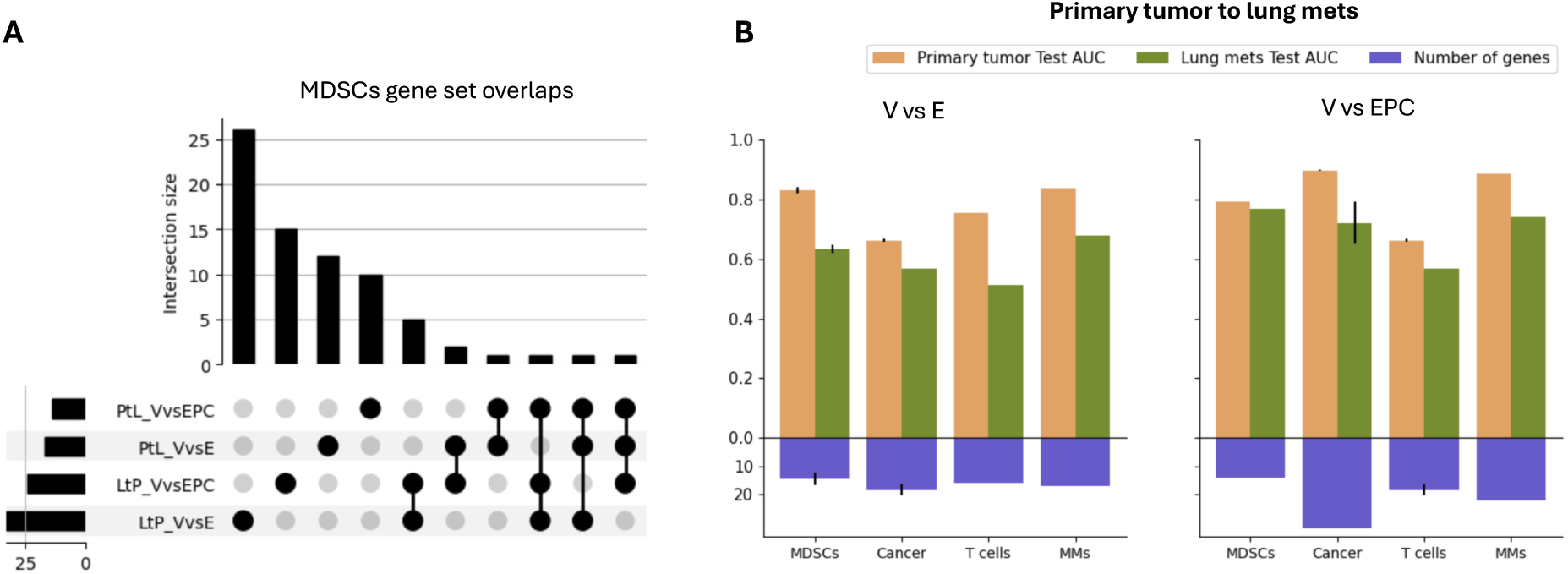
Application of iLR on lung mets and primary tumor. (**A**) Upset plot showing the overlap among the iLR gene set from the four comparison of MDSCs: lung mets V vs E, lung mets V vs EPC, primary tumor V vs E and primary tumor V vs EPC. (**B**) The test AUC for lung mets and primary tumor using the gene set draw from the primary tumor.

**Figure S4:**
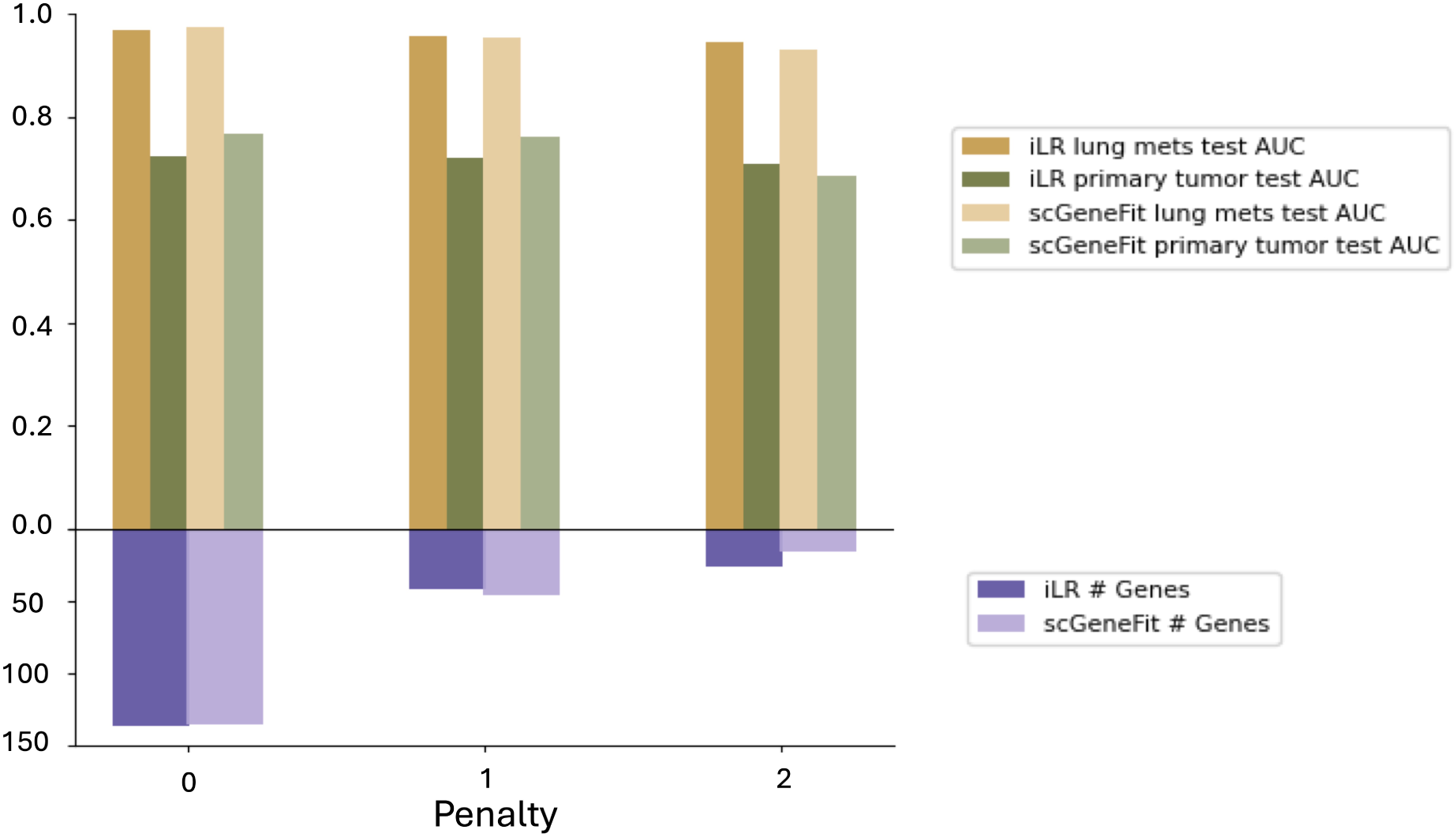
Comparison of classification AUC of iLR and scGeneFit on MDSCs V vs. **E.** Comparing the classification test AUC on lung mets and test AUC on primary tumor with the genes identified from lung mets with the number of genes on the bottom.

**Figure S5:**
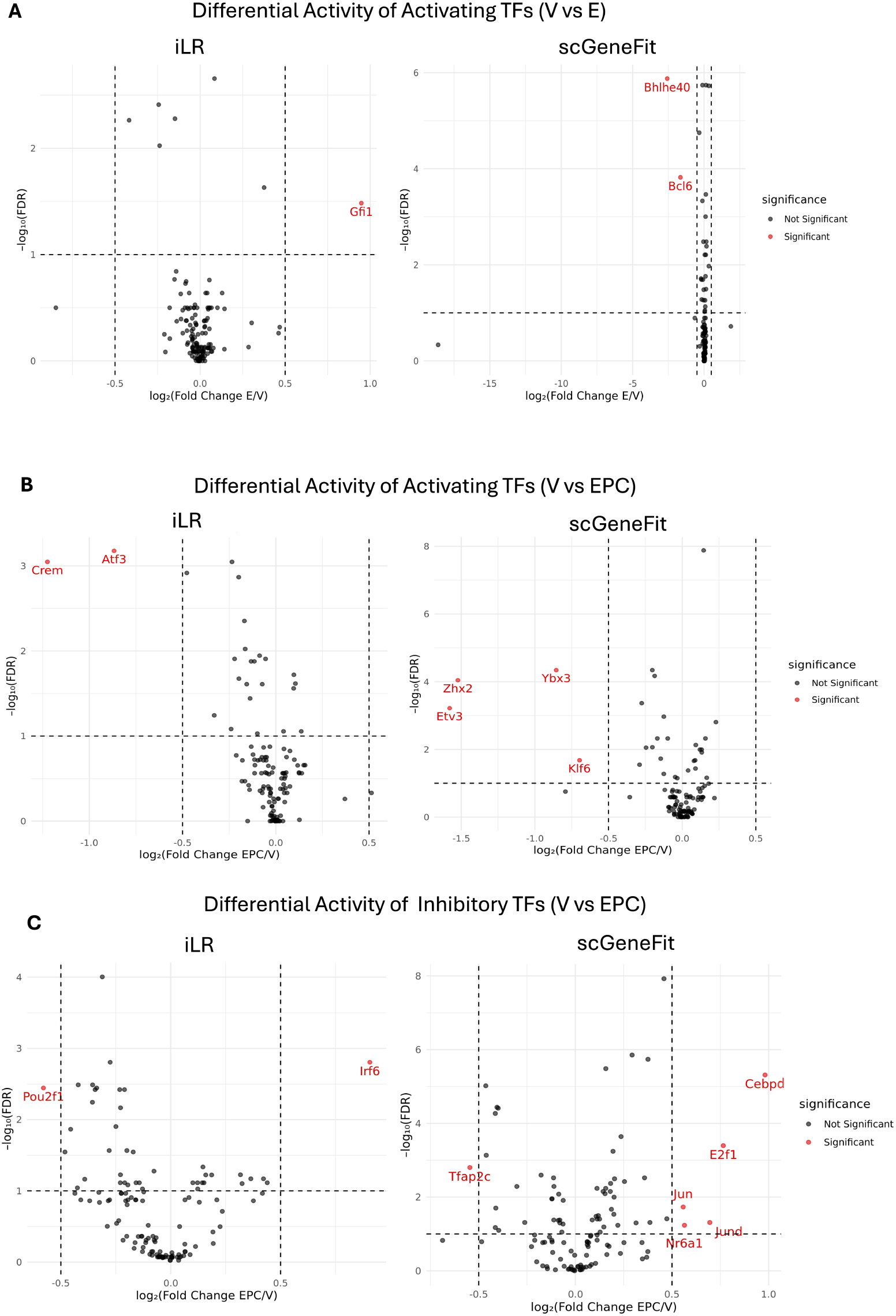
TFs significantly changed inferred by SCORPION from iLR and scGeneFit genes in MDSCs. (**A**) Significantly differential activity of activating TFs comparing V and E for iLR and scGeneFit genes respectively. (**B**)Significantly differential activity of activating TFs comparing V and EPC for iLR and scGeneFit genes respectively. (**C**)Significantly differential activity of inhibitory TFs comparing V and EPC for iLR and scGeneFit genes respectively.

## References

Abrahams, B. S., Arking, D. E., Campbell, D. B., Mefford, H. C., Morrow, E. M., Weiss, L. A., Menashe, I., Wadkins, T., Banerjee-Basu, S., & Packer, A. (2013). SFARI gene 2.0: A community-driven knowledgebase for the autism spectrum disorders (ASDs). Molecular Autism, 4, 36. 10.1186/2040-2392-4-36

Anders, S., & Huber, W. (2010). Differential expression analysis for sequence count data. Genome Biology, 11(10), R106. 10.1186/gb-2010-11-10-r106

Badia-i-Mompel, P., Vélez Santiago, J., Braunger, J., Geiss, C., Dimitrov, D., Müller-Dott, S., Taus, P., Dugourd, A., Holland, C. H., Ramirez Flores, R. O., & Saez-Rodriguez, J. (2022). decoupleR: Ensemble of computational methods to infer biological activities from omics data. Bioinformatics Advances, 2(1), vbac016. 10.1093/bioadv/vbac016

Baugh, A. G., Gonzalez, E., Narumi, V. H., Kreger, J., Liu, Y., Rafie, C., Castanon, S., Jang, J., Kagohara, L. T., Anastasiadou, D. P., Leatherman, J., Armstrong, T., Chan, I., Karagiannis, G. S., Jaffee, E. M., MacLean, A., & Roussos Torres, E. T. (2024). A new neu—a syngeneic model of spontaneously metastatic HER2-positive breast cancer. Clinical & Experimental Metastasis, 41(5), 733–746. 10.1007/s10585-024-10289-z

Baugh, A. G., Gonzalez, E., Narumi, V. H., Kreger, J., Liu, Y., Rafie, C., Castanon, S., Jang, J., Kagohara, L. T., Anastasiadou, D. P., Leatherman, J., Armstrong, T., Chan, I., Karagiannis, G. S., Jaffee, E. M., MacLean, A., & Torres, E. T. R. (2024). A new Neu—a syngeneic model of spontaneously metastatic HER2-positive breast cancer. Clinical & Experimental Metastasis. 10.1007/s10585-024-10289-z

Berkson, J. (1944). Application of the Logistic Function to Bio-Assay. Journal of the American Statistical Association, 39(227), 357–365. 10.1080/01621459.1944.10500699

Chea, S., Kreger, J., Lopez-Burks, M. E., MacLean, A. L., Lander, A. D., & Calof, A. (2023). Gastrulation-stage gene expression in Nipbl+/- mouse embryos foreshadows the development of syndromic birth defects. 10.1101/2023.10.16.558465

Christmas, B. J., Rafie, C. I., Hopkins, A. C., Scott, B. A., Ma, H. S., Cruz, K. A., Woolman, S., Armstrong, T. D., Connolly, R. M., Azad, N. A., Jaffee, E. M., & Roussos Torres, E. T. (2018). Entinostat Converts Immune-Resistant Breast and Pancreatic Cancers into Checkpoint-Responsive Tumors by Reprogramming Tumor-Infiltrating MDSCs. Cancer Immunology Research, 6(12), 1561–1577. 10.1158/2326-6066.CIR-18-0070

Cox, D. R. (1958). The Regression Analysis of Binary Sequences. Journal of the Royal Statistical Society. Series B (Methodological), 20(2), 215–242.

Crowell, H. L., Morillo Leonardo, S. X., Soneson, C., & Robinson, M. D. (2023). The shaky foundations of simulating single-cell RNA sequencing data. Genome Biology, 24(1), 1–19. 10.1186/s13059-023-02904-1

Dawson, M. A., & Kouzarides, T. (2012). Cancer epigenetics: From mechanism to therapy [Publisher: Elsevier]. Cell, 150(1), 12–27. 10.1016/j.cell.2012.06.013

Dumitrascu, B., Villar, S., Mixon, D. G., & Engelhardt, B. E. (2021). Optimal marker gene selection for cell type discrimination in single cell analyses [Publisher: Nature Publishing Group]. Nature Communications, 12(1), 1186. 10.1038/s41467-021-21453-4

El Khoury, W., & Nasr, Z. (2021). Deregulation of ribosomal proteins in human cancers. Bioscience Reports, 41(12), BSR20211577. 10.1042/BSR20211577

Falkenberg, K. J., & Johnstone, R. W. (2014). Histone deacetylases and their inhibitors in cancer, neurological diseases and immune disorders [Publisher: Nature Publishing Group]. Nature Reviews Drug Discovery, 13(9), 673–691. 10.1038/nrd4360

Fang, Z., Liu, X., & Peltz, G. (2023). GSEApy: A comprehensive package for performing gene set enrichment analysis in python. Bioinformatics, 39(1), btac757. 10.1093/bioinformatics/btac757

Feuermann, M., Mi, H., Gaudet, P., Muruganujan, A., Lewis, S. E., Ebert, D., Mushayahama, T., & Thomas, P. D. (2025). A compendium of human gene functions derived from evolutionary modelling. Nature, 1–9. 10.1038/s41586-025-08592-0

Fitzgerald, J. B., Schoeberl, B., Nielsen, U. B., & Sorger, P. K. (2006). Systems biology and combination therapy in the quest for clinical efficacy [Publisher: Nature Publishing Group]. Nature Chemical Biology, 2(9), 458–466. 10.1038/nchembio817

Garcia-Alonso, L., Holland, C. H., Ibrahim, M. M., Turei, D., & Saez-Rodriguez, J. (2019). Benchmark and integration of resources for the estimation of human transcription factor activities. Genome Research, 29(8), 1363–1375. 10.1101/gr.240663.118

Hu, C., Yang, K., Li, M., Huang, W., Zhang, F., & Wang, H. (2018). Lipocalin 2: A potential therapeutic target for breast cancer metastasis [Publisher: Dove Medical Press _eprint: https://www.tandfonline.com/doi/pdf/10.2147/OTT.S181223]. OncoTargets and Therapy, 11, 8099–8106. 10.2147/OTT.S181223

Huang, Y., & Zhang, P. (2021). Evaluation of machine learning approaches for cell-type identification from single-cell transcriptomics data. Briefings in Bioinformatics, 22(5), bbab035. 10.1093/bib/bbab035

Jiao, X., Nawab, O., Patel, T., Kossenkov, A. V., Halama, N., Jaeger, D., & Pestell, R. G. (2019). Recent advances targeting CCR5 for cancer and its role in immuno-oncology. Cancer Research, 79(19), 4801–4807. 10.1158/0008-5472.CAN-19-1167

Jin, S., Guerrero-Juarez, C. F., Zhang, L., Chang, I., Ramos, R., Kuan, C.-H., Myung, P., Plikus, M. V., & Nie, Q. (2021). Inference and analysis of cell-cell communication using CellChat. Nature Communications, 12(1), 1088. 10.1038/s41467-021-21246-9

Kamimoto, K., Stringa, B., Hoffmann, C. M., Jindal, K., Solnica-Krezel, L., & Morris, S. A. (2023). Dissecting cell identity via network inference and in silico gene perturbation. Nature, 1–10. 10.1038/s41586-022-05688-9

Kreger, J., Roussos Torres, E. T., & MacLean, A. L. (2023). Myeloid-Derived Suppressor–Cell Dynamics Control Outcomes in the Metastatic Niche. Cancer Immunology Research, OF1–OF15. 10.1158/2326-6066.CIR-22-0617

Le, H., Peng, B., Uy, J., Carrillo, D., Zhang, Y., Aevermann, B. D., & Scheuermann, R. H. (2022). Machine learning for cell type classification from single nucleus RNA sequencing data [Publisher: Public Library of Science]. PLOS ONE, 17(9), e0275070. 10.1371/journal.pone.0275070

Lopez, R., Regier, J., Cole, M. B., Jordan, M. I., & Yosef, N. (2018). Deep generative modeling for single-cell transcriptomics. Nature Methods, 15(12), 1053–1058. 10.1038/s41592-018-0229-2

Ma, Y.-F., Chen, Y., Fang, D., Huang, Q., Luo, Z., Qin, Q., Lin, J., Zou, C., Huang, M., Meng, D., Huang, Q., & Lu, G.-M. (2021). The immune-related gene CD52 is a favorable biomarker for breast cancer prognosis. Gland Surgery, 10(2), 780–798. 10.21037/ gs-20-922

Mitchell, J. T., Stapleton, O., Krishnan, K., Nagaraj, S., Lvovs, D., Cherry, C., Poissonnier, A., Horton, W., Adey, A., Rao, V., Huff, A., Zimmerman, J. W., Kagohara, L. T., Zaidi, N., Coussens, L. M., Jaffee, E. M., Elisseeff, J. H., & Fertig, E. J. (2025). Differential cell signaling testing for cell-cell communication inference from single-cell data by dominoSignal. 10.1101/2025.05.02.651747

Mitra, R., & MacLean, A. L. (2021). RVAgene: Generative modeling of gene expression time series data. Bioinformatics, (btab260). 10.1093/bioinformatics/btab260

Müller-Dott, S., Tsirvouli, E., Vázquez, M., Flores, R. O. R., Badia-i-Mompel, P., Fallegger, R., Lægreid, A., & Saez-Rodriguez, J. (2023, April 1). Expanding the coverage of regulons from high-confidence prior knowledge for accurate estimation of transcription factor activities [Pages: 2023.03.30.534849 Section: New Results]. 10.1101/ 2023.03.30.534849

Nelson, M. E., Riva, S. G., & Cvejic, A. (2022). SMaSH: A scalable, general marker gene identification framework for single-cell RNA-sequencing. BMC Bioinformatics, 23(1), 328. 10.1186/s12859-022-04860-2

Osorio, D., Capasso, A., Eckhardt, S. G., Giri, U., Somma, A., Pitts, T. M., Lieu, C. H., Messersmith, W. A., Bagby, S. M., Singh, H., Das, J., Sahni, N., Yi, S. S., & Kuijjer, M. L. (2024a). Population-level comparisons of gene regulatory networks modeled on high-throughput single-cell transcriptomics data. Nature Computational Science, 4(3), 237–250. 10.1038/s43588-024-00597-5

Osorio, D., Capasso, A., Eckhardt, S. G., Giri, U., Somma, A., Pitts, T. M., Lieu, C. H., Messersmith, W. A., Bagby, S. M., Singh, H., Das, J., Sahni, N., Yi, S. S., & Kuijjer, M. L. (2024b). Population-level comparisons of gene regulatory networks modeled on high-throughput single-cell transcriptomics data [Publisher: Nature Publishing Group]. Nature Computational Science, 4(3), 237–250. 10.1038/s43588-024-00597-5

Pedregosa, F., Varoquaux, G., Gramfort, A., Michel, V., Thirion, B., Grisel, O., Blondel, M., Prettenhofer, P., Weiss, R., Dubourg, V., Vanderplas, J., Passos, A., Cournapeau, D., Brucher, M., Perrot, M., & Duchesnay, É. (2011). Scikit-learn: Machine learning in python. Journal of Machine Learning Research, 12(85), 2825–2830. http://jmlr.org/ papers/v12/pedregosa11a.html

Perri, F., Longo, F., Giuliano, M., Sabbatino, F., Favia, G., Ionna, F., Addeo, R., Della Vittoria Scarpati, G., Di Lorenzo, G., & Pisconti, S. (2017). Epigenetic control of gene expression: Potential implications for cancer treatment. Critical Reviews in Oncology/Hematology, 111, 166–172. 10.1016/j.critrevonc.2017.01.020

Pullin, J. M., & McCarthy, D. J. (2024). A comparison of marker gene selection methods for single-cell RNA sequencing data. Genome Biology, 25(1), 56. 10.1186/s13059-024-03183-0

Qiu, X., Zhou, T., Li, S., Wu, J., Tang, J., Ma, G., Yang, S., Hu, J., Wang, K., Shen, S., Wang, H., & Chen, L. (2024). Spatial single-cell protein landscape reveals vimentinhigh macrophages as immune-suppressive in the microenvironment of hepatocellular carcinoma [Publisher: Nature Publishing Group]. Nature Cancer, 5(10), 1557–1578. 10.1038/ s43018-024-00824-y

Robinson, M. D., McCarthy, D. J., & Smyth, G. K. (2010). edgeR: A bioconductor package for differential expression analysis of digital gene expression data. Bioinformatics, 26(1), 139–140. 10.1093/bioinformatics/btp616

Rommelfanger, M. K., Behrends, M., Chen, Y., Martinez, J., Bens, M., Xiong, L., Rudolph, K. L., & MacLean, A. L. (2023). Gene regulatory network inference with popInfer reveals dynamic regulation of hematopoietic stem cell quiescence upon diet restriction and aging. 10.1101/2023.04.18.537360

Roussos Torres, E. T., Ho, W. J., Danilova, L., Tandurella, J. A., Leatherman, J., Rafie, C., Wang, C., Brufsky, A., LoRusso, P., Chung, V., Yuan, Y., Downs, M., O’Connor, A., Shin, S. M., Hernandez, A., Engle, E. L., Piekarz, R., Streicher, H., Talebi, Z., . . . Connolly, R. M. (2024). Entinostat, nivolumab and ipilimumab for women with advanced HER2-negative breast cancer: A phase Ib trial. Nature Cancer, 1–14. 10.1038/s43018-024-00729-w

Sidiropoulos, D. N., Rafie, C. I., Jang, J. K., Castanon, S., Baugh, A. G., Gonzalez, E., Christmas, B. J., Narumi, V. H., Davis-Marcisak, E. F., Sharma, G., Bigelow, E., Vaghasia, A., Gupta, A., Skaist, A., Considine, M., Wheelan, S. J., Ganesan, S. K., Yu, M., Yegnasubramanian, S., . . . Roussos Torres, E. T. (2022). Entinostat decreases immune suppression to promote antitumor responses in a HER2+ breast tumor microenvironment. Cancer Immunology Research, 10(5), 656–669. 10.1158/2326-6066.CIR-21-0170

Sun, Y., & Qiu, P. (2024). Hierarchical marker genes selection in scRNA-seq analysis [Publisher: Public Library of Science]. PLOS Computational Biology, 20(12), e1012643. 10.1371/journal.pcbi.1012643

Svensson, V., da Veiga Beltrame, E., & Pachter, L. (2020). A curated database reveals trends in single-cell transcriptomics. Database, 2020, baaa073. 10.1093/database/baaa073

Szklarczyk, D., Kirsch, R., Koutrouli, M., Nastou, K., Mehryary, F., Hachilif, R., Gable, A. L., Fang, T., Doncheva, N. T., Pyysalo, S., Bork, P., Jensen, L. J., & von Mering, C. (2023). The STRING database in 2023: Protein-protein association networks and functional enrichment analyses for any sequenced genome of interest. Nucleic Acids Research, 51, D638–D646. 10.1093/nar/gkac1000

Vargo, A. H. S., & Gilbert, A. C. (2020). A rank-based marker selection method for high throughput scRNA-seq data. BMC Bioinformatics, 21(1), 477. 10.1186/ s12859-020-03641-z

Velmeshev, D., Schirmer, L., Jung, D., Haeussler, M., Perez, Y., Mayer, S., Bhaduri, A., Goyal, N., Rowitch, D. H., & Kriegstein, A. R. (2019). Single-cell genomics identifies cell type–specific molecular changes in autism. Science, 364(6441), 685–689. 10.1126/science.aav8130

Wamsley, B., Bicks, L., Cheng, Y., Kawaguchi, R., Quintero, D., Margolis, M., Grundman, J., Liu, J., Xiao, S., Hawken, N., Mazariegos, S., & Geschwind, D. H. (2024). Molecular cascades and cell type–specific signatures in ASD revealed by single-cell genomics. Science, 384(6698), eadh2602. 10.1126/science.adh2602

Wang, L., Trasanidis, N., Wu, T., Dong, G., Hu, M., Bauer, D. E., & Pinello, L. (2023). Dictys: Dynamic gene regulatory network dissects developmental continuum with single-cell multiomics. Nature Methods, 1–11. 10.1038/s41592-023-01971-3

Wilcoxon, F. (1945). Individual Comparisons by Ranking Methods. Biometrics Bulletin, 1(6), 80–83.

Wolf, F. A., Angerer, P., & Theis, F. J. (2018). SCANPY: Large-scale single-cell gene expression data analysis. Genome Biology, 19(1), 15. 10.1186/s13059-017-1382-0

Yan, L., Cheng, J., Nie, Q., & Sun, X. (2025). Dissecting multilayer cell-cell communications with signaling feedback loops from spatial transcriptomics data. Genome Research. 10.1101/gr.279857.124

Zappia, L., Phipson, B., & Oshlack, A. (2017). Splatter: Simulation of single-cell RNA sequencing data. Genome Biology, 18(1), 174. 10.1186/s13059-017-1305-0

