## Supplementary Tables and Figures for "Inference of marker genes of subtle cell state changes via iterative logistic regression"

### Supplementary Tables and Figures for
### Inference of marker genes of subtle cell state changes via iterative logistic regression

Yingtong Liu[1], Aaron G. Baugh[2], Evanthia T. Roussos Torres[2,*], & Adam L. MacLean[1,*]
[1]Department of Quantitative and Computational Biology, University of Southern California, Los Angeles, CA 90089, USA

[2]Department of Medicine, Division of Medical Oncology, Keck School of Medicine, Norris Comprehensive Cancer Center, University of Southern California, Los Angeles, CA 90033, USA

| DE scale factor | Sample size | Number of DE genes | Number of iLR genes (penalty = 1) |
|---|---|---|---|
| 0.1 | 500 | 4 | 16 |
| 0.1 | 2000 | 34 | 16 |
| 0.2 | 500 | 28 | 16 |
| 0.2 | 2000 | 85 | 20 |
| 0.25 | 500 | 42 | 12 |
| 0.25 | 2000 | 105 | 20 |
| 0.5 | 500 | 81 | 16 |
| 1 | 500 | 222 | 12 |

**S1 Table.** The number of significant differentially expressed genes (adjusted p-value cutoff 0.05) identified by Wilcoxon rank sum test in eight simulated data.

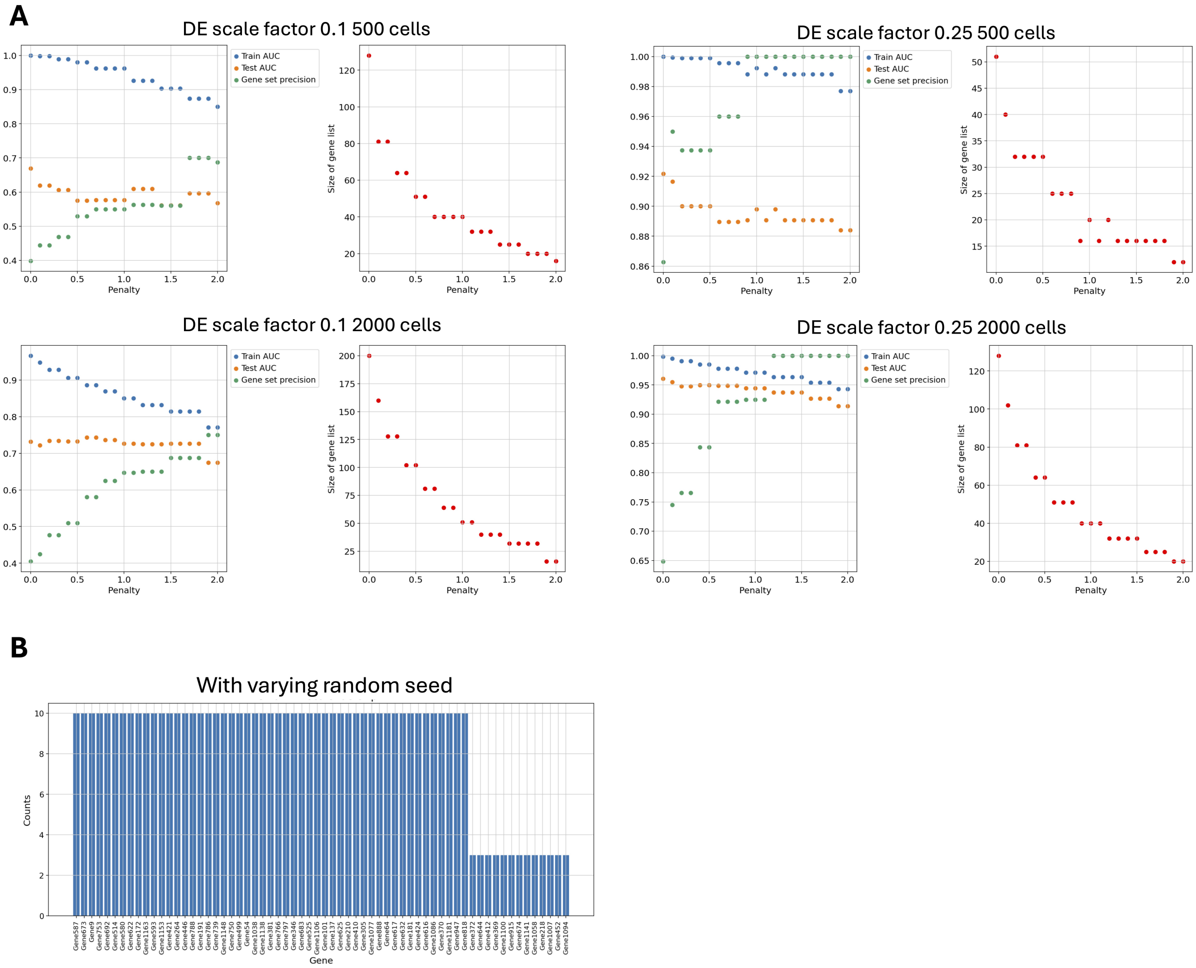


**S1 Figure. iLR evaluation on simulated data.**(**A**) Scatter plots trace the penalty effect on four simulated data with different DE scale factor or sample size. (**B**) Examination the iLR robustness by running 10 times, varying the random seed.

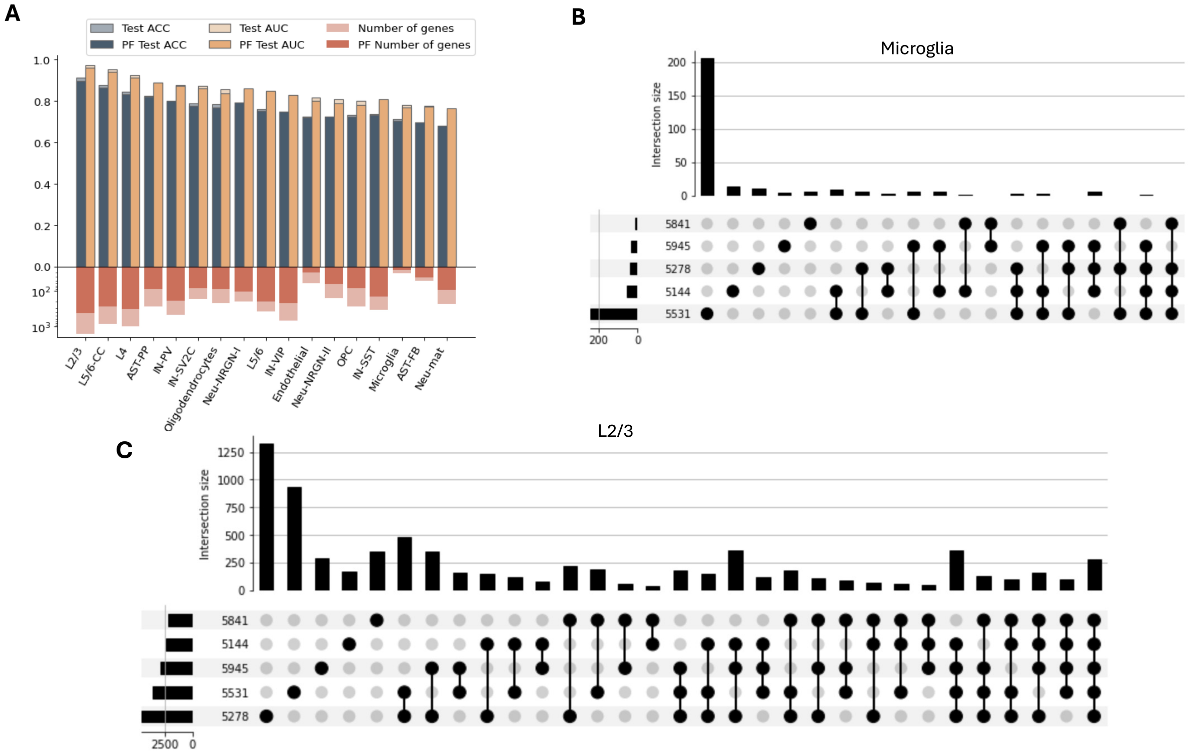


**S2 Figure. iLR application on ASD dataset.**(**A**) iLR classification accuracy and gene set size comparing no pareto front (solid colored) and with pareto front without penalty (pale colored). (**B**) Upset plot of Wilcoxon rank sum test gene sets identified by comparison of microglia of individual patient and all controls. (**C**) Upset plot of Wilcoxon rank sum test gene sets identified by comparison of L2/3 of individual patient and all controls.

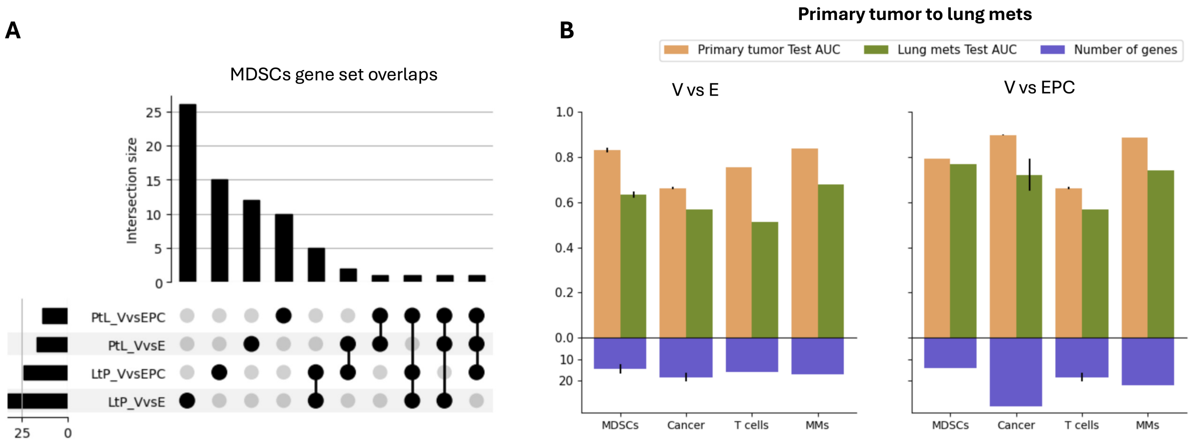


**S3 Figure. Application of iLR on lung mets and primary tumor.**(**A**) Upset plot showing the overlap among the iLR gene set from the four comparison of MDSCs: lung mets V vs E, lung mets V vs EPC, primary tumor V vs E and primary tumor V vs EPC. (**B**) The test AUC for lung mets and primary tumor using the gene set draw from the primary tumor.

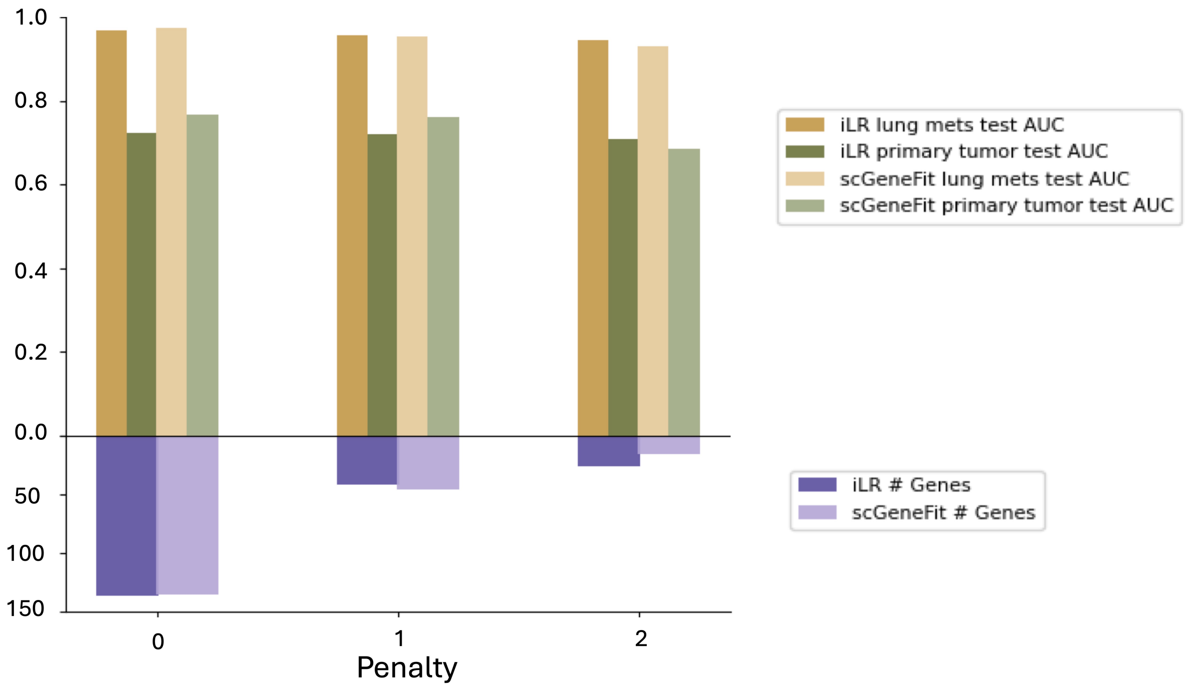


**S4 Figure. Comparison of classification AUC of iLR and scGeneFit on MDSCs V vs E.** Comparing the classification test AUC on lung mets and test AUC on primary tumor with the genes identified from lung mets with the number of genes on the bottom.

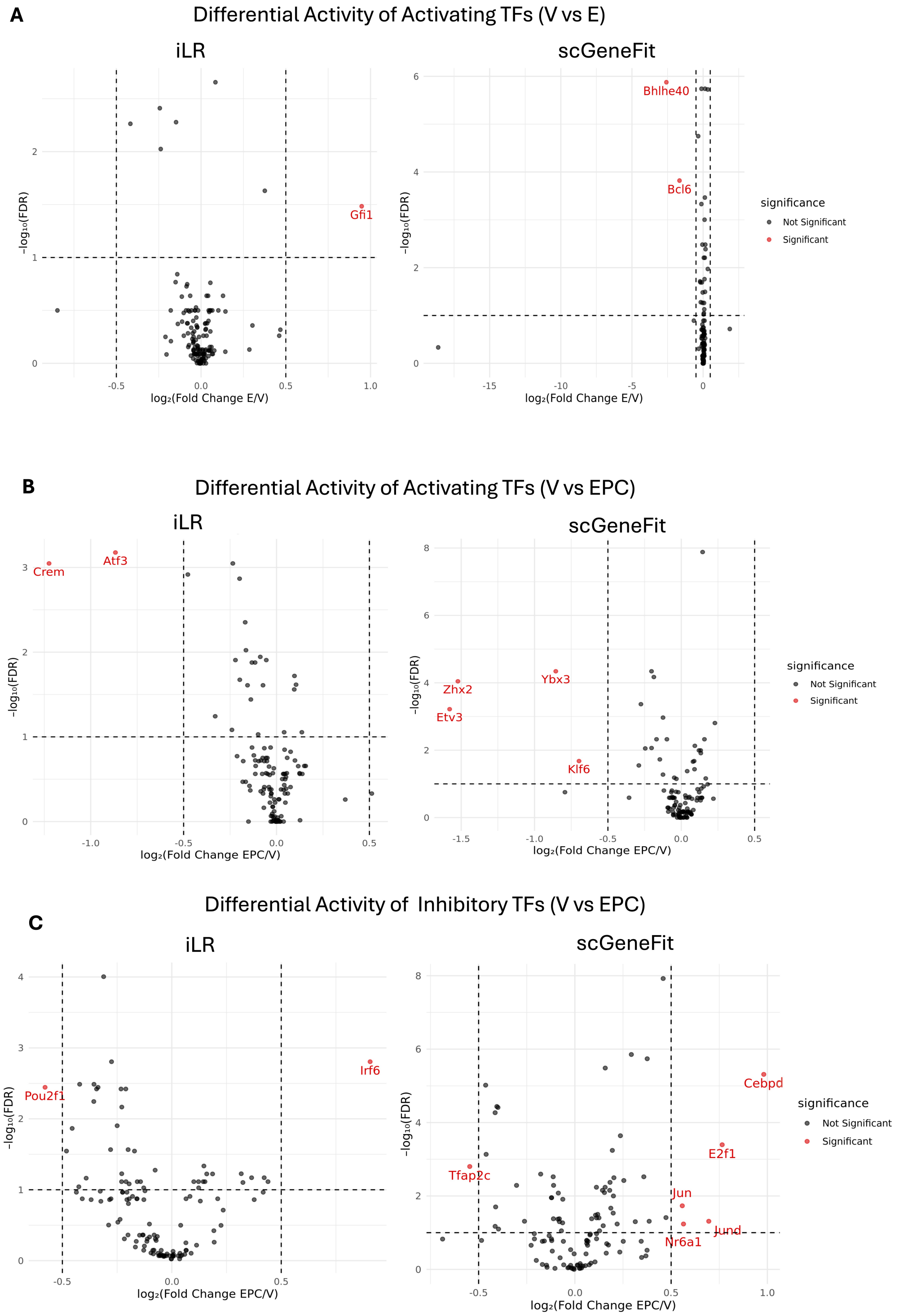


**S5 Figure. TFs significantly changed inferred by SCORPION from iLR and scGeneFit genes in MDSCs (A)** Significantly differential activity of activating TFs comparing V and E for iLR and scGeneFit genes respectively. (**B**)Significantly differential activity of activating TFs comparing V and EPC for iLR and scGeneFit genes respectively. (**C**)Significantly differential activity of inhibitory TFs comparing V and EPC for iLR and scGeneFit genes respectively.
